# Pre-existing antibiotic tolerance facilitates plasmid-mediated carbapenem resistance evolution in clinical *Klebsiella pneumoniae*

**DOI:** 10.64898/2026.08.24.746623

**Authors:** Wei-Li Zhang, Bo Zheng, Meng-Lan Zhou, Rong Zhang, Ying-Chun Xu, Jia-Feng Liu

## Abstract

Antibiotic tolerance enables bacteria to survive bactericidal antibiotic exposure and has been linked to resistance evolution in laboratory systems and individual infections, but its role in plasmid-mediated resistance evolution in clinical populations remains unclear. Here, we analyzed a longitudinal collection of > 800 clinical *Klebsiella pneumoniae* isolates spanning 1997–2020. Among 779 minimum inhibitory concentration (MIC)-defined ertapenem-susceptible isolates, 137 (17.6%) displayed hidden ertapenem tolerance, defined by enhanced survival after 6 h at 30 × isolate-specific ertapenem MIC, mostly without extended lag time or reduced growth rate. Tolerance was detected before local ertapenem introduction and was enriched among ertapenem-resistant isolates, supporting a population-level association between pre-existing tolerance and the emergence of carbapenem resistance. Genomic and plasmid-curing analyses separated plasmid-mediated carbapenem resistance from plasmid-independent antibiotic tolerance. Moreover, tolerant recipient backgrounds enhanced resistance plasmid acquisition, preserved viable recipients following antibiotic exposure and accelerated ceftazidime–avibactam resistance evolution. A phylogeny-guided variant-enrichment analysis further identified the *uhpABC* regulatory operon as a candidate tolerance-associated locus, and coordinated expression of the complete operon increased ertapenem survival. Together, these findings identify clinical antibiotic tolerance as a pre-existing, MIC-hidden phenotype that can facilitate plasmid-mediated carbapenem resistance evolution in *K. pneumoniae*.

## Introduction

Antibiotic resistance poses a major global health threat, and Gram-negative pathogens such as *Klebsiella pneumoniae* (*K. pneumoniae*) present a particular therapeutic challenge^1,2^. As a major cause of hospital-acquired infections, *K. pneumoniae* has become increasingly difficult to treat because of the global expansion of multidrug-resistant lineages, particularly carbapenem-resistant clones^3,4^. Thus, carbapenem-resistant *K. pneumoniae* (CRKP) is recognized by the World Health Organization (WHO) as a priority bacterial pathogen^5^. The emergence and spread of CRKP are facilitated by the horizontal acquisition and dissemination of carbapenemase-encoding plasmids^6–8^. However, clinical surveillance and treatment decisions rely primarily on minimum inhibitory concentration (MIC)-based susceptibility classification. Because MIC-based testing measures growth inhibition rather than bacterial survival during antibiotic exposure^9^, clinically relevant survival heterogeneity may remain undetected by routine susceptibility testing^10–12^. Consequently, the prevalence and temporal distribution of antibiotic tolerance in clinical *K. pneumoniae* populations remain poorly defined. Unlike resistance, which enables bacterial growth at elevated antibiotic concentrations, tolerance enables bacteria to survive transient antibiotic exposure without necessarily increasing the MIC, and has been implicated in treatment failure and infection recurrence^13–15^. Experimental studies have shown that tolerance can evolve rapidly under repeated antibiotic exposure^16–18^ and can facilitate the subsequent evolution of resistance under laboratory conditions^19,20^. More recently, longitudinal tracking of individual infections suggested that tolerance can also shape resistance evolution within the host^21–24^. However, these studies have largely focused on laboratory evolution or individual infection trajectories, in which resistance often arises through chromosomal mutation. Together, these observations raise the question of whether antibiotic tolerance can shape plasmid-mediated resistance evolution in clinical *K. pneumoniae* populations.

To address this question, we analyzed a longitudinal clinical collection of *K. pneumoniae* isolates by integrating MIC profiling, antibiotic survival phenotyping, comparative genomics and functional assays. We found that antibiotic tolerance exists as a hidden, pre-existing phenotype within MIC-defined susceptible populations. This phenotype was enriched among ertapenem-resistant isolates but was separable from resistance plasmid carriage. Tolerant recipient backgrounds enhanced resistance-plasmid acquisition and preserved viable recipients during antibiotic exposure, thereby promoting plasmid-mediated resistance evolution. These findings identify clinical antibiotic tolerance as a pre-existing, regulated survival trait that can shape the emergence and dissemination of carbapenem resistance in *K. pneumoniae*.

## Results

### Hidden ertapenem tolerance in MIC-defined ertapenem-susceptible isolates is largely uncoupled from extended lag or slow growth

To investigate the evolutionary relationship between ertapenem tolerance and resistance, we assembled a longitudinal collection of clinical *K. pneumoniae* isolates from Peking Union Medical College Hospital, China, spanning 1997–2020, a period encompassing years before and after ertapenem was introduced into clinical use at this hospital in 2006 (Extended Data Fig. 1a). Ertapenem MICs, determined by broth microdilution, were used to stratify isolates into ertapenem-susceptible and ertapenem-resistant groups. The MIC distribution shifted progressively upward over time, with high-level ertapenem-resistant isolates (MIC > 100 μg/mL) detected predominantly in later years (Fig. 1a), consistent with the global expansion of CRKP^25,7^. However, because MICs measure growth inhibition rather than survival after bactericidal antibiotic exposure, these data did not determine whether tolerant phenotypes were already present within the MIC-defined susceptible population.

**Fig. 1.**
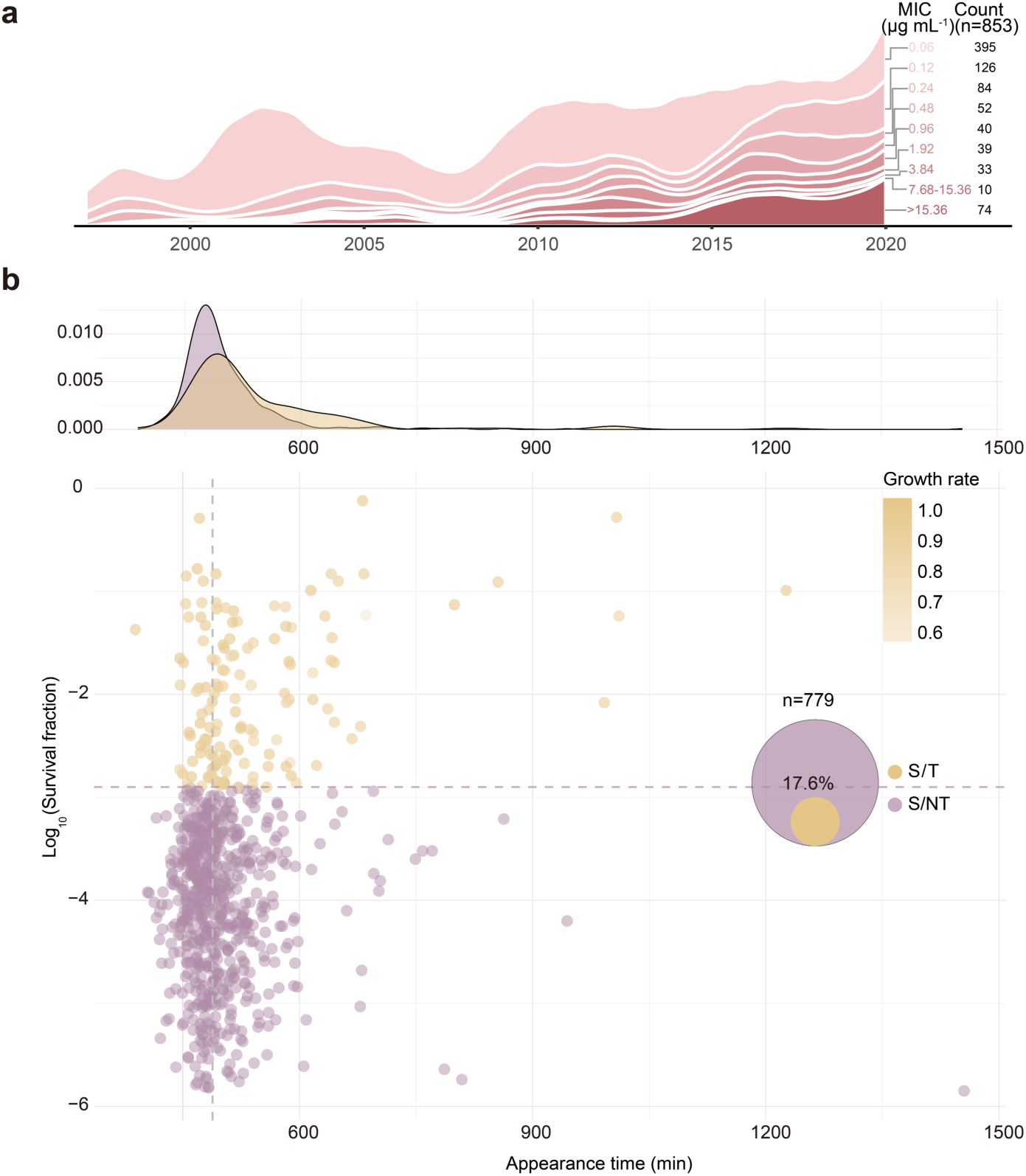
Longitudinal ertapenem MIC profiling reveals hidden tolerance among MIC-defined susceptible *Klebsiella pneumoniae*. **a**, Longitudinal distribution of ertapenem MICs among clinical *K. pneumoniae* isolates collected between 1997 and 2020 (n = 853). The stacked distribution plot shows the temporal distribution of isolates across ertapenem MIC categories, with the total number of isolates in each category indicated on the right. For visualization, the two isolates with an ertapenem MIC of 15.36 μg ml^-1^ were combined with the 7.68 μg ml^-1^ category and are shown as the 7.68–15.36 μg ml^-1^ bin. Among the 74 isolates with ertapenem MICs >15.36 μg ml^-1^, 70 had MICs > 100 μg ml^-1^, whereas the remaining four had MICs between 15.36 and 100 μg ml^-1^. **b**, Survival–appearance-time landscape of MIC-defined ertapenem-susceptible isolates (n = 779). Each point represents one isolate, plotted according to mean colony appearance time measured by ScanLag and survival after 6 h of exposure to ertapenem at 30× the isolate-specific MIC. Isolates showing a ≥ 10-fold increase in survival relative to *K. pneumoniae* ATCC 43816 were classified as susceptible/tolerant (S/T; yellow, n = 137), whereas the remaining isolates were classified as susceptible/non-tolerant (S/NT; purple, n = 642). The horizontal dashed line indicates the operational tolerance threshold, and the vertical dashed line indicates the mean colony appearance time of *K. pneumoniae* ATCC 43816 measured under the same ScanLag conditions. The top density plot shows the distribution of colony appearance times. Among S/T isolates, color intensity indicates growth rate. The inset shows the prevalence of hidden ertapenem tolerance within the MIC-defined ertapenem-susceptible population (17.6%, 137/779).

We therefore established a standardized killing assay to quantify ertapenem tolerance among MIC-defined ertapenem-susceptible isolates (Extended Data Fig. 1b). To account for MIC heterogeneity across the cohort, each isolate was exposed to ertapenem at 30 × its respective MIC (Extended Data Fig. 1c). Based on time-kill kinetics, survival after 6 h of exposure was selected as the operational readout for tolerance (Extended Data Fig. 1c). Isolates showing a ≥ 10-fold increase in survival relative to the reference strain ATCC 43816 were classified as tolerant, enabling stratification of ertapenem-susceptible isolates into non-tolerant (S/NT) and tolerant (S/T) groups. Overall, 17.6% (137/779) of MIC-defined ertapenem-susceptible isolates exceeded this threshold (Fig. 1b), revealing a hidden tolerant subset within a population that would otherwise be classified as susceptible by MIC testing alone.

Previous studies, largely based on laboratory-evolved models, have identified two canonical forms of antibiotic tolerance^14^, termed “tolerance by lag” and “tolerance by slow growth”, in which extended lag phase or reduced growth rate enhances survival during antibiotic exposure. We therefore asked whether the clinical S/T isolates identified here conformed to this canonical framework. Using ScanLag, we quantified colony appearance-time distributions as a proxy for lag-phase duration across the cohort. However, most S/T isolates (114/137, 83.2%) displayed colony appearance-time distributions comparable to those of S/NT isolates and the reference strain ATCC 43816, and did not meet our operational criterion for “tolerance by lag”, defined as an appearance-time delay of ≥ 600 min relative to the reference strain (Fig. 1b). Likewise, exponential growth rates were comparable between S/T isolates and the reference strain, indicating that most S/T isolates lacked a measurable growth-rate defect (Fig. 1b).

Together, these findings reveal that MIC-defined susceptible clinical *K. pneumoniae* isolates include a substantial ertapenem-tolerant subset that is not captured by MIC-based classification. Notably, most of these tolerant isolates did not exhibit extended lag phase or reduced growth rate, distinguishing this clinical ertapenem tolerance from the extended-lag or slow-growth tolerance commonly described in laboratory-evolved models.

### Pre-existing ertapenem tolerance may reflect broader β-lactam-associated survival

We next mapped the longitudinal distribution of S/T isolates across the clinical collection. Although ertapenem was introduced into clinical practice at Peking Union Medical College Hospital in 2006, S/T isolates were already detectable between 1997 and 2005 (Fig. 2a). This temporal pattern suggests that ertapenem tolerance was unlikely to have arisen solely as a direct response to local ertapenem use.

**Fig. 2.**
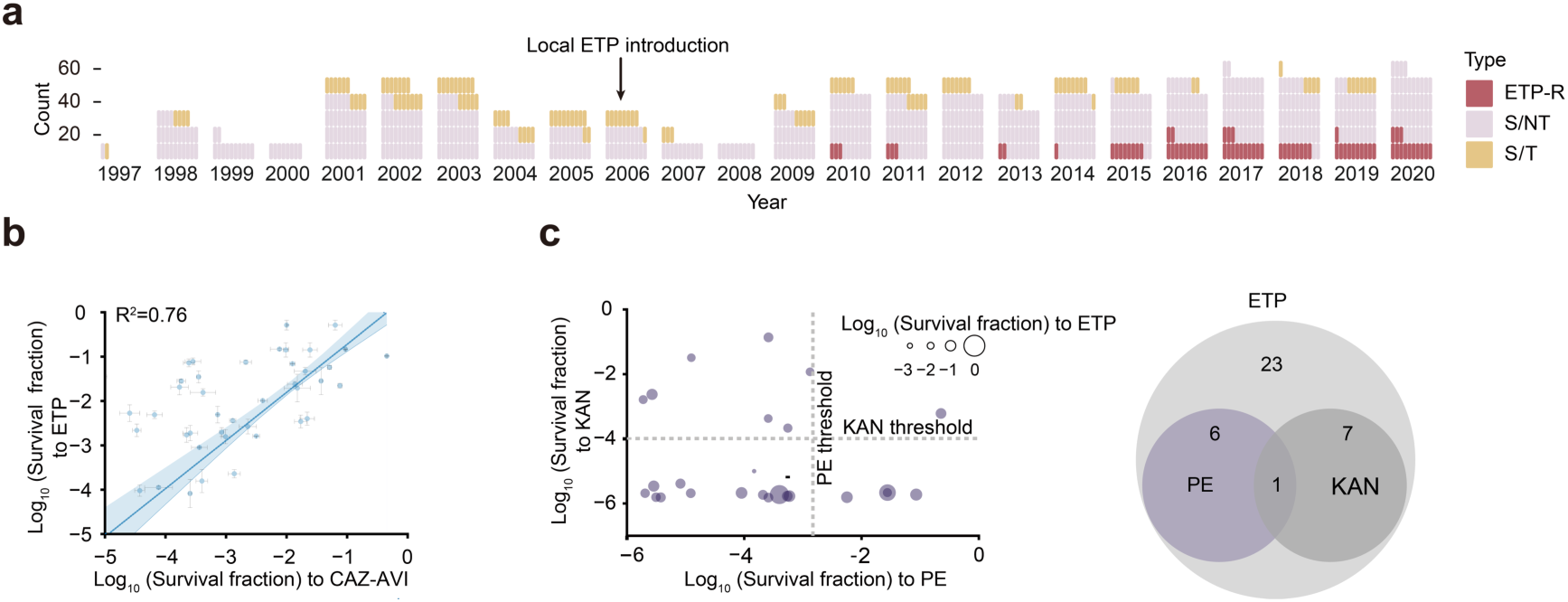
Pre-existing ertapenem tolerance is associated with β-lactam survival and limited non-β-lactam cross-tolerance. **a**, Annual distribution of clinical *K. pneumoniae* isolates classified as ertapenem-resistant (ETP-R; red), MIC-defined ertapenem-susceptible/non-tolerant (S/NT; purple) or MIC-defined ertapenem-susceptible/tolerant (S/T; yellow). Each vertical stack represents isolates collected in the indicated year. The arrow indicates the local introduction of ertapenem in 2006. **b**, Correlation between survival under ertapenem (ETP) and ceftazidime–avibactam (CAZ–AVI) exposure among MIC-defined ertapenem-susceptible isolates (n = 43). Each point represents one clinical isolate, plotted according to the mean log₁₀-transformed survival fraction after 6 h of exposure to each antibiotic at 30 × the isolate-specific MIC. Error bars indicate s.d. from biological replicates. The fitted line and shaded region indicate the linear regression and 95% confidence interval; R^2^ is shown. **c**, Cross-antibiotic survival among MIC-defined ertapenem-susceptible/tolerant (S/T) isolates tested under polymyxin E (PE) and kanamycin (KAN) exposure. Each point represents one isolate, plotted according to the mean log_10_-transformed survival fraction under PE and KAN exposure. Point size indicates the mean survival fraction under ETP exposure. Dashed lines indicate the operational tolerance thresholds for PE and KAN. The Venn diagram summarizes the overlap among isolates classified as tolerant to ETP, PE and KAN, with numbers indicating isolate counts in each subset.

Previous studies have shown that antibiotic tolerance can confer cross-protection during exposure to different antibiotics, including β-lactams^17,11^. We therefore asked whether this pre-existing tolerance reflected broader β-lactam-associated survival. Because many isolates in this clinical collection were already resistant to older β-lactams, including earlier-generation cephalosporins, these drugs were unsuitable for standardized killing assays to quantify tolerance. To test this possibility, we examined survival under ceftazidime–avibactam (CAZ– AVI) exposure, a β-lactam/β-lactamase inhibitor combination used as a last-line treatment option for carbapenem-resistant *K. pneumoniae* infections^27^. Across ertapenem-susceptible isolates spanning different tolerance levels, survival under CAZ–AVI positively correlated with survival under ertapenem (Fig. 2b), linking ertapenem tolerance to elevated survival in a distinct β-lactam treatment context.

To determine whether this ertapenem tolerance was restricted to β-lactam exposure, we further tested survival under non-β-lactam antibiotics, including kanamycin and polymyxin E. Although cross-class survival was not universal, a subset of isolates displayed elevated survival across multiple antibiotic classes (Fig. 2c). Thus, pre-existing ertapenem tolerance was most directly associated with β-lactam-related survival, whereas in some isolates elevated survival extended to non-β-lactam antibiotics.

### Tolerance is enriched during ertapenem resistance evolution but is not directly conferred by resistance plasmid carriage

Previous experimental evolution studies have proposed a stepwise adaptive trajectory in which antibiotic tolerance arises before resistance and facilitates the emergence of partial resistance, which can ultimately progress to high-level resistance under sustained antibiotic selection^28^. In our longitudinal clinical collection, S/T isolates were already detectable before ertapenem was introduced at this hospital, whereas ertapenem-resistant isolates were detected predominantly after ertapenem use began (Fig. 2a), revealing a population-level temporal pattern consistent with this experimentally proposed trajectory. If similar evolutionary trajectories occur in clinical populations, tolerant phenotypes might remain detectable and become enriched among ertapenem-resistant isolates. We therefore quantified the prevalence of tolerant phenotypes in this ertapenem-resistant population.

Direct quantification of ertapenem tolerance in ertapenem-resistant isolates is technically challenging because their ertapenem MICs frequently exceed 100 μg/mL. We therefore turned to CAZ–AVI survival as a surrogate readout for tolerance in this group. This approach was technically feasible because most ertapenem-resistant isolates remained susceptible to CAZ– AVI (66/74, 89.2%; MIC ≤ 8 μg/mL) (Fig. 3a), enabling quantitative survival measurements under bactericidal conditions. This choice was further supported by the positive correlation between CAZ–AVI and ertapenem survival observed above in MIC-defined ertapenem-susceptible isolates (Fig. 2b). Using this CAZ–AVI-based readout, ertapenem-resistant isolates were stratified into ertapenem-resistant non-tolerant (R/NT) and ertapenem-resistant tolerant (R/T) groups (Extended Data Fig. 2a).

**Fig. 3.**
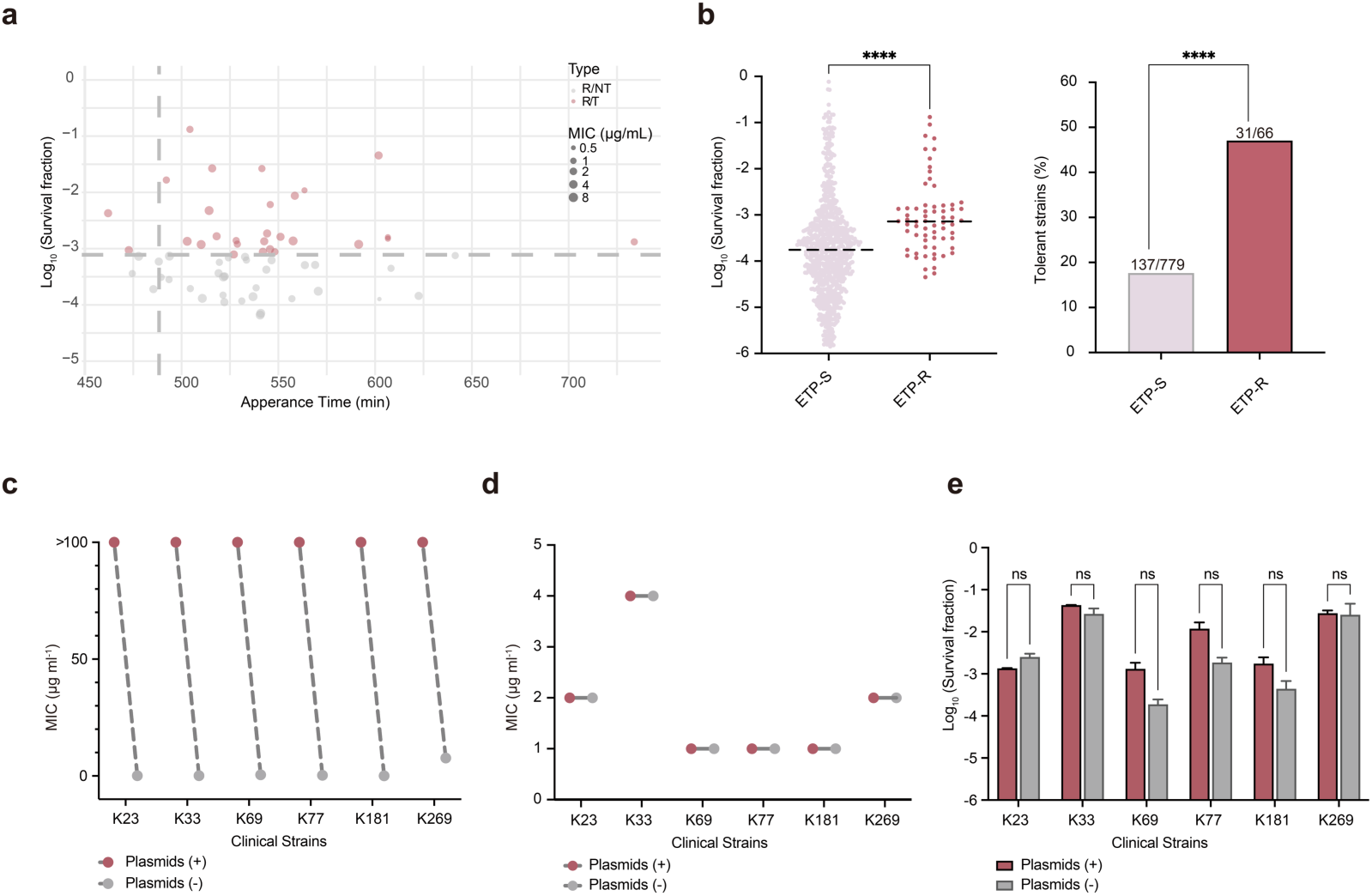
Tolerance is enriched among ertapenem-resistant isolates and separable from resistance plasmid carriage. **a**, Ceftazidime–avibactam survival, appearance-time and MIC landscape of ertapenem-resistant *K. pneumoniae* isolates. Each point represents one isolate, plotted according to mean colony appearance time and log_10_-transformed survival fraction after 6 h of exposure to CAZ–AVI at 30× the isolate-specific MIC. Point size indicates CAZ–AVI MIC. Resistant/non-tolerant (R/NT) and resistant/tolerant (R/T) isolates are shown in grey and red, respectively. The horizontal dashed line indicates the operational tolerance threshold, and the vertical dashed line indicates the mean colony appearance time of *K. pneumoniae* ATCC 43816 measured under the same ScanLag conditions. **b**, Enrichment of tolerance among ertapenem-resistant isolates. Left, distribution of survival fractions among MIC-defined ertapenem-susceptible and ertapenem-resistant isolates. Survival was assessed by ertapenem killing assays for MIC-defined ertapenem-susceptible isolates and by CAZ–AVI killing assays for ertapenem-resistant isolates suitable for CAZ–AVI survival profiling. Right, proportion of tolerant isolates in each group, showing enrichment of tolerance among ertapenem-resistant isolates: 17.6% (137/779) versus 47.0% (31/66). Numbers above bars indicate tolerant isolates relative to the total number of isolates tested. **c**, **d**, MICs of representative *bla*_KPC-2_-positive clinical isolates before and after plasmid curing. Ertapenem MICs are shown in c, and CAZ– AVI MICs are shown in d. Plasmid-positive parental isolates and plasmid-cured derivatives are shown in red and grey, respectively. **e**, CAZ–AVI survival of parental and plasmid-cured derivatives. Bars show log_10_-transformed survival fractions after CAZ–AVI exposure. Data are presented as mean ± s.d. from at least three biological replicates. Statistical significance was determined using a two-sided Mann–Whitney U test for **b**, left, a two-sided Fisher’s exact test for **b**, right, and two-way ANOVA followed by Šídák’s multiple-comparisons test for **e**. \*\*\*\**P* < 0.0001; ns, not significant.

Among the assayable ertapenem-resistant isolates, 47.0% (31/66) were classified as R/T (Fig. 3b), representing a 2.7-fold enrichment relative to the frequency of S/T isolates in the MIC-defined ertapenem-susceptible population (17.6%, 137/779). Although tolerance was not universally retained across resistant isolates, this significant enrichment is consistent with clinically resistant populations preferentially emerging from tolerant backgrounds. As observed for S/T isolates, R/T isolates did not show a marked colony appearance-time delay (Fig. 3a). Using an analogous survival assay, tolerance was also detected among NDM-producing CRKP isolates (15.7%, 8/51) (Extended Data Fig. 2b), suggesting that this survival phenotype was not confined to KPC-producing resistant isolates.

The enrichment of tolerance among resistant isolates raised an important mechanistic question. Because carbapenem resistance in clinical *K. pneumoniae* is often mediated by horizontally acquired resistance plasmids rather than by chromosomal mutations alone^29,30^ , the observed association between tolerance and resistance could reflect either pre-existing tolerance facilitating resistance plasmid acquisition or resistance plasmids themselves directly enhancing survival under antibiotic exposure.

To distinguish between these possibilities, we first characterized the genetic basis of carbapenem resistance within the clinical collection. Consistent with previous epidemiological studies^31,32^, *bla*_KPC-2_ was the dominant carbapenemase gene among sequenced isolates with high-level ertapenem resistance (64/68, 94.1%; Extended Data Table 1), whereas the remaining four isolates carried *bla*_NDM_-type carbapenemases. MOB-suite-based plasmid reconstruction further showed that all detected carbapenemase genes were located on plasmid-associated contigs, suggesting that high-level ertapenem resistance in the sequenced subset was largely plasmid mediated.

To determine whether resistance plasmids directly contributed to the tolerant phenotype, representative KPC-2-producing isolates were subjected to plumbagin-mediated plasmid curing^33^ (Extended Data Fig. 3a). This treatment reduced ertapenem MICs by approximately 2–4 orders of magnitude (Fig. 3c), confirming that high-level ertapenem resistance was primarily plasmid encoded. In contrast, CAZ–AVI MICs (Fig. 3d) and survival under CAZ–AVI exposure (Fig. 3e) remained largely unchanged after plasmid curing. Colony appearance dynamics also showed only minor alterations (Extended Data Fig. 3b), further supporting a plasmid-independent tolerant phenotype.

Collectively, these findings indicate that tolerance is enriched during ertapenem resistance evolution but is not directly conferred by resistance plasmid carriage. Combined with the observation that tolerance preceded detectable ertapenem resistance, these results suggest that tolerance may represent a pre-existing phenotypic background for plasmid-mediated carbapenem resistance emergence in this clinical cohort.

### Tolerant backgrounds facilitate plasmid-mediated resistance evolution by enhancing plasmid acquisition and amplifying transfer through recipient survival

Because conjugative plasmid transfer is a major route for the spread of carbapenem resistance in clinical *K. pneumoniae*, we asked whether tolerant backgrounds facilitate resistance plasmid acquisition and transfer. To minimize potential confounding by bacterial chromosomal background, we reconstructed a recombination-filtered core-genome SNP phylogeny of representative clinical isolates. Most CRKP isolates belonged to a closely related lineage (58/69, 84.1%) (Fig. 4a and Extended Data Fig. 4), enabling comparisons of conjugative plasmid acquisition within a relatively homogeneous lineage. Within this closely related resistant lineage, R/T isolates carried *bla*_KPC-2_-bearing plasmids assigned to multiple plasmid clusters (Extended Data Fig. 5). Although one plasmid cluster was predominant, R/T isolates were not confined to a single plasmid background. Together with their close core-genome relatedness, this pattern was consistent with repeated acquisition of resistance plasmids within tolerant backgrounds rather than expansion solely following a single plasmid-acquisition event. We therefore compared conjugative plasmid acquisition in closely related R/T and R/NT isolates, using BW25113/p3R-4 carrying *bla*_NDM_ as the donor strain. R/T isolates exhibited an approximately tenfold higher conjugation efficiency than R/NT isolates (Fig. 4b). Thus, even in the absence of antibiotic selection during conjugation, tolerant recipient backgrounds were associated with enhanced acquisition of resistance plasmids.

**Fig. 4.**
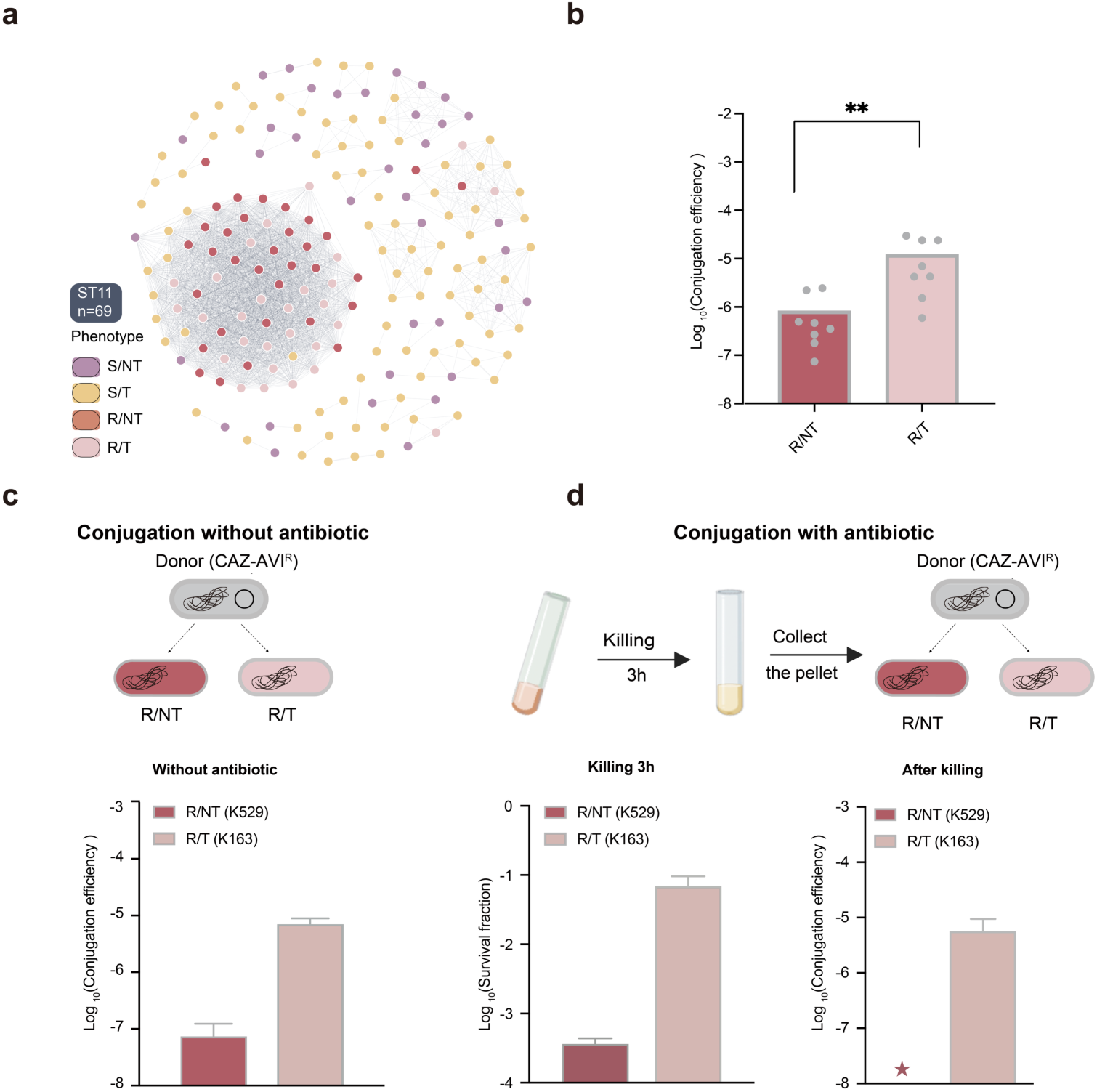
Tolerant recipient backgrounds enhance resistance plasmid acquisition and preserve conjugative transfer after ceftazidime–avibactam exposure. **a**, Core-genome-based cluster network of sequenced clinical *K. pneumoniae* isolates, visualized using Gephi v0.10. Each node represents one isolate, and node color indicates phenotypic category: MIC-defined ertapenem-susceptible non-tolerant (S/NT), MIC-defined ertapenem-susceptible tolerant (S/T), ertapenem-resistant non-tolerant (R/NT) or ertapenem-resistant tolerant (R/T). Grey lines connect isolate pairs defined as genetically related in the core-genome relatedness network. The dominant ST11 cluster is indicated. **b**, Baseline conjugation efficiency of R/NT and R/T recipient isolates in the absence of antibiotic exposure. Each group included eight independent recipient isolates. Bars indicate group means, and points represent isolate-level mean conjugation efficiencies calculated from three biological replicates. **c**, Schematic and quantification of conjugation assays performed without CAZ–AVI exposure, using representative R/NT (K529) and R/T (K163) recipients and a CAZ–AVI-resistant donor. Bars indicate log10-transformed conjugation efficiency for the indicated recipient strains. **d**, Schematic and quantification of conjugation assays performed after 3 h CAZ–AVI exposure. Representative R/NT (K529) and R/T (K163) recipients were exposed to CAZ–AVI for 3 h before mating; surviving cells were collected and used as recipients in conjugation assays with a CAZ– AVI-resistant donor. Left, survival after CAZ–AVI exposure. Right, conjugation efficiency among surviving recipients after CAZ–AVI exposure. The red asterisk indicates no detectable transconjugants. For assays **c** and **d**, data are presented as mean ± s.d. from at least three biological replicates. Statistical comparison in **b** was performed using isolate-level mean conjugation efficiencies and a two-sided Mann–Whitney U test. \*\**P* < 0.01.

We next asked whether antibiotic exposure further amplified this plasmid-acquisition advantage of tolerant backgrounds. Representative R/T and R/NT isolates were subjected to antibiotic treatment before conjugation assays (Fig. 4c). After antibiotic exposure, R/T recipients exhibited approximately 100-fold greater survival than R/NT recipients (Fig. 4d). Conjugative transfer remained readily detectable in surviving R/T recipients but was nearly abolished in R/NT recipients (Fig. 4d). Thus, antibiotic-mediated killing magnified pre-existing differences in plasmid acquisition by maintaining a larger pool of viable tolerant recipients capable of conjugative plasmid transfer.

Together, these findings suggest that tolerant backgrounds promote plasmid-mediated resistance evolution not only by enhancing baseline plasmid acquisition but also by amplifying transfer through increased recipient survival following antibiotic exposure.

### Pre-existing tolerance predicts accelerated evolution of CAZ–AVI resistance

Having established a retrospective association between tolerance and resistance in the clinical collection, we next asked whether pre-existing tolerance could predict the subsequent evolution of CAZ–AVI resistance. To model the transition from established ertapenem resistance to subsequent CAZ–AVI selection, we selected four ertapenem-resistant clinical isolates that remained susceptible to CAZ–AVI but differed in CAZ–AVI tolerance, comprising three R/T isolates and one R/NT isolate.

Consistent with their tolerance classification, R/T isolates exhibited pronounced survival advantages relative to the R/NT isolate following transient CAZ–AVI exposure (Fig. 5a). We then established an in vitro evolution framework to model recurrent antibiotic exposure, consisting of repeated cycles of CAZ–AVI killing, recovery, and regrowth (Fig. 5b). Replicate populations derived from R/T isolates evolved CAZ–AVI resistance significantly faster than those derived from the R/NT isolate (Fig. 5c). During serial passaging, R/NT-derived populations maintained relatively stable MICs, whereas R/T-derived populations rapidly evolved high-level CAZ–AVI resistance within a few evolutionary cycles.

**Fig. 5.**
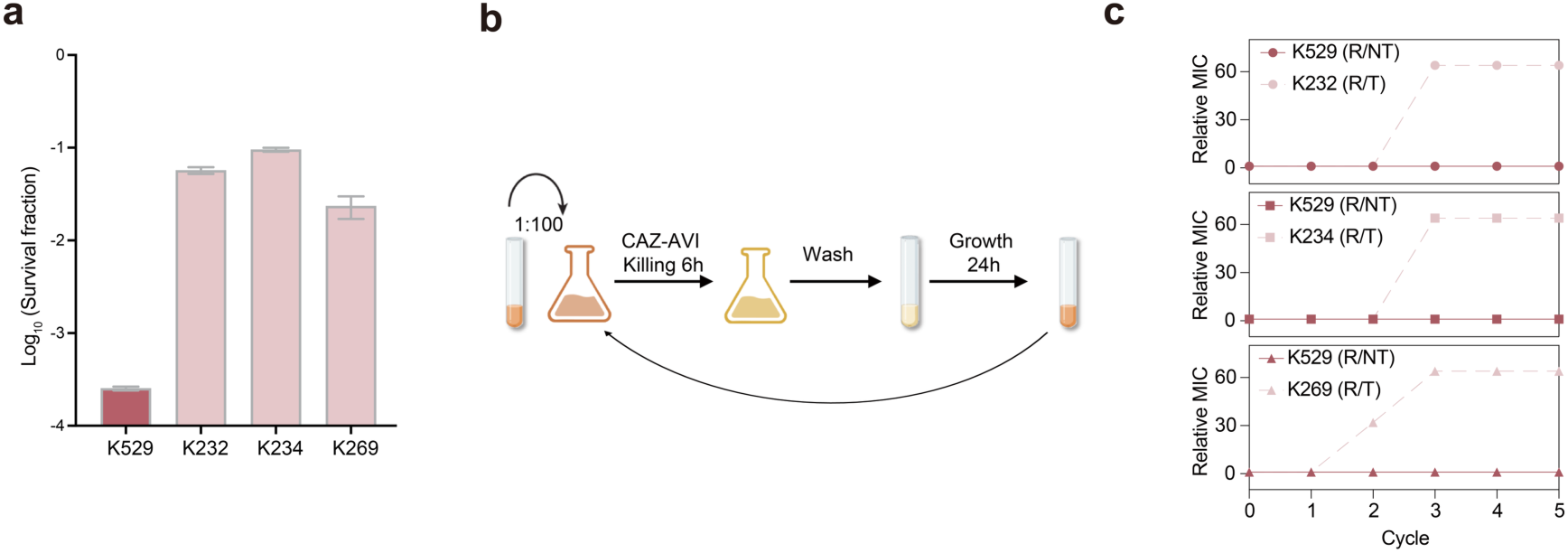
Pre-existing tolerance is associated with accelerated evolution of ceftazidime– avibactam resistance. **a**, Ceftazidime–avibactam (CAZ–AVI) survival of ertapenem-resistant clinical isolates selected for in vitro evolution. The ertapenem-resistant non-tolerant isolate K529 (R/NT) and three ertapenem-resistant tolerant isolates, K232, K234 and K269 (R/T), were exposed to CAZ–AVI for 6 h at 30× the isolate-specific MIC. Bars show log_10_-transformed survival fractions; data are presented as mean ± s.d. from three biological replicates. **b**, Schematic of the in vitro evolution assay used to model recurrent CAZ–AVI exposure. Cultures were diluted 1:100, exposed to CAZ–AVI for 6 h, washed and regrown for 24 h before the next cycle. **c**, Evolution of CAZ–AVI MICs during repeated antibiotic exposure. Relative MIC values were calculated by normalizing the MIC at each cycle to the corresponding starting MIC at cycle 0; thus, cycle 0 is set to 1 for each lineage. Solid lines indicate the R/NT-derived lineage from K529, and dashed lines indicate R/T-derived lineages from K232, K234 and K269. The K529-derived R/NT lineage remained largely stable, whereas R/T-derived lineages rapidly evolved elevated CAZ–AVI MICs.

These results indicate that pre-existing tolerance can facilitate the subsequent evolution of CAZ–AVI resistance under repeated antibiotic exposure. Together with our retrospective analyses, these findings position tolerance as both a contributor to plasmid-mediated resistance emergence and a candidate predictor of subsequent resistance evolution.

### Clinical variation and functional validation implicate the *uhpABC* operon in ertapenem tolerance

To identify genetic determinants associated with antibiotic tolerance in clinical isolates while minimizing confounding from broad population structure, we used the core-genome phylogeny to select a closely related lineage in which isolates differed in ertapenem tolerance despite limited core-genome divergence (Fig. 6a). Within this lineage, variant-enrichment analysis identified the *uhpABC* locus as a candidate tolerance-associated region, with the strongest signal mapping to *uhpB* and exceeding the FDR threshold (Fig. 6b). A complementary cohort-level frequency comparison further showed that the leading *uhpB* variant was enriched, although not exclusive, among tolerant isolates (Extended Data Fig. 6), suggesting that this signal was not confined to the discovery lineage.

**Fig. 6.**
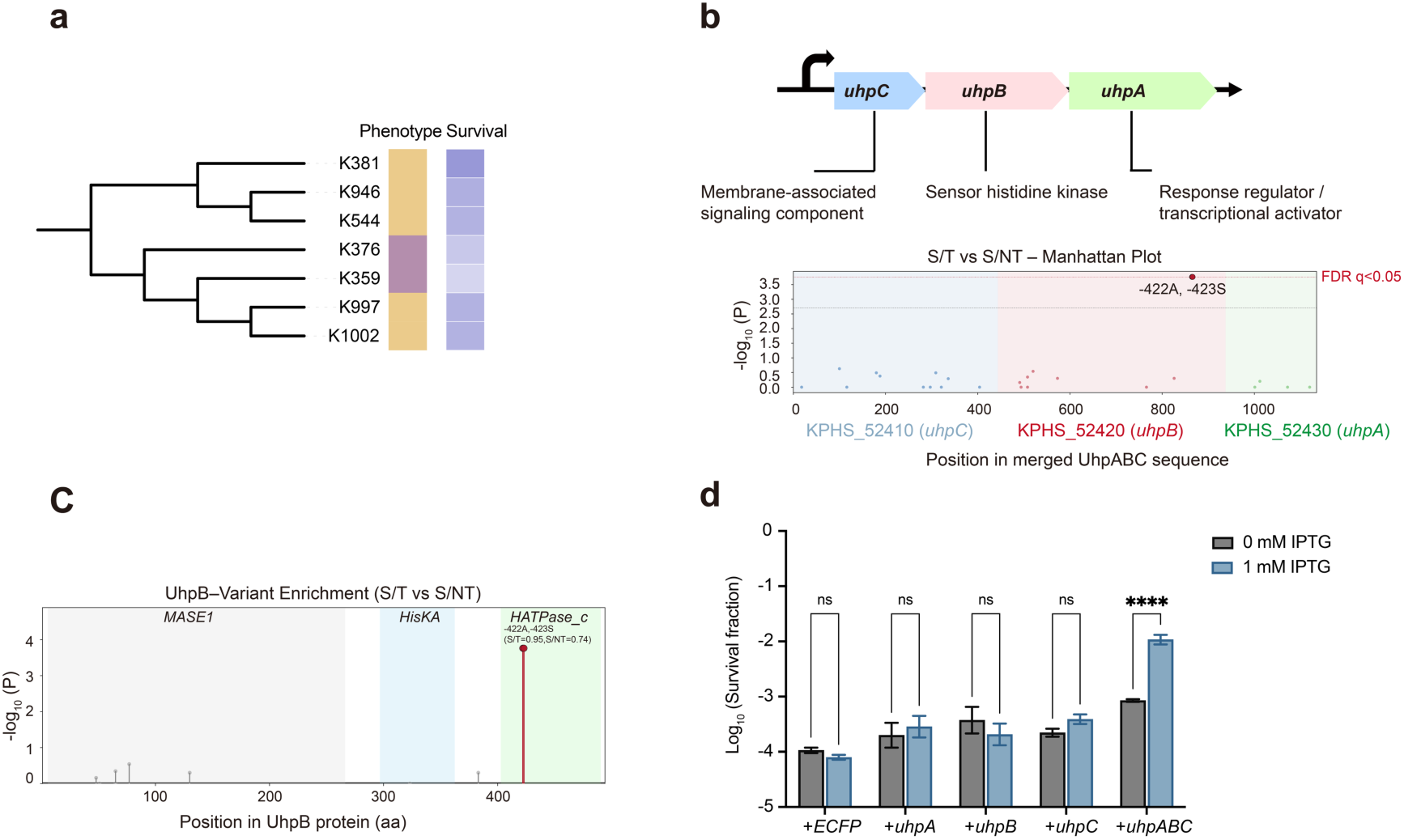
Phylogeny-guided variant analysis identifies *uhpABC* as a candidate tolerance-associated operon. **a**, Zoomed view of a phylogenetically clustered focal lineage selected from the core-genome phylogeny for within-lineage comparative analysis. This lineage contained both ertapenem-susceptible tolerant isolates (S/T; yellow) and ertapenem-susceptible non-tolerant isolates (S/NT; purple) despite limited core-genome divergence. The annotation bar indicates tolerance status, and the adjacent heat map shows isolate-level ertapenem survival, with darker shades indicating higher survival. **b**, Schematic of the *uhpABC* locus and variant-enrichment analysis across the merged *uhpABC* region. The locus encodes a putative signal-transduction module comprising *UhpA*, a response regulator and transcriptional activator; *UhpB*, a predicted sensor histidine kinase; and *UhpC*, a membrane-associated signaling component. The Manhattan-style plot shows variant enrichment between tolerant and non-tolerant isolates within the focal lineage. The strongest enrichment signal mapped to *uhpB* and exceeded the FDR q < 0.05 threshold. **c**, Mapping of tolerance-enriched variants onto the predicted *UhpB* domain architecture. The leading uhpB variant mapped to the C-terminal HATPase_c domain, a region associated with histidine kinase activity. **d**, Ertapenem survival of *K. pneumoniae* ATCC 43816 carrying IPTG-inducible expression constructs for individual *uhp* genes or the complete *uhpABC* operon. Strains carrying the empty vector control (+ECFP), *uhpA*, *uhpB*, *uhpC* or *uhpABC* were cultured in the absence or presence of 1 mM IPTG and exposed to ertapenem. Induction of *uhpA*, *uhpB* or *uhpC* alone did not significantly increase survival, whereas induction of the complete *uhpABC* operon significantly enhanced ertapenem survival. Bars show log_10_-transformed survival fractions; data are presented as mean ± s.d. from three biological replicates. Statistical significance was determined by two-way ANOVA followed by Šídák’s multiple-comparisons test. \*\*\*\**P* < 0.0001; ns, not significant.

The *uhpABC* locus encodes a predicted signal-transduction module comprising UhpB, a sensor histidine kinase; UhpA, a response regulator; and UhpC, a membrane-associated signaling component. This organization is consistent with the canonical UhpABC system in enterobacteria, which senses extracellular glucose-6-phosphate and regulates UhpT-dependent hexose phosphate uptake^34^. Protein-domain mapping localized the enriched *uhpB* variant to the C-terminal HATPase_c domain of UhpB, a region associated with histidine kinase activity (Fig. 6c). Together, the lineage-based enrichment, cohort-level distribution, and domain localization nominated *uhpB*, and more broadly the *uhpABC* regulatory system, as candidate components associated with clinical ertapenem tolerance.

We next tested whether the UhpABC system could functionally modulate antibiotic survival. IPTG-inducible constructs expressing individual *uhp* components or the complete *uhpABC* operon were introduced into *K. pneumoniae* ATCC 43816. Induced expression of *uhpA*, *uhpB*, or *uhpC* alone did not significantly increase survival under ertapenem exposure. In contrast, coordinated expression of the complete *uhpABC* operon increased survival by approximately one order of magnitude relative to the vector control (Fig. 6d). These results indicate that operon-level activation of the UhpABC system is sufficient to enhance ertapenem survival in a reference *K. pneumoniae* background.

## Discussion

Our study extends previous mutation-centered models linking antibiotic tolerance to resistance evolution by showing that, in clinical *K. pneumoniae*, tolerance can also contribute to plasmid-mediated carbapenem resistance evolution. Tolerance was detectable among MIC-defined ertapenem-susceptible isolates before high-level ertapenem resistance became established and was enriched among ertapenem-resistant isolates, supporting a population-level association between pre-existing tolerance and carbapenem resistance emergence. Because carbapenem resistance in this cohort was largely plasmid mediated, these findings suggest that tolerant recipient backgrounds can shape resistance evolution not only through survival under antibiotic exposure, but also by influencing resistance plasmid acquisition and selection.

A central implication of our work is that MIC-based susceptibility testing captures only one dimension of antibiotic response. MIC assays measure growth inhibition, whereas tolerance reflects bacterial survival during bactericidal antibiotic exposure^11,12^. By phenotyping a longitudinal clinical collection spanning the local introduction of ertapenem, we identified a tolerant subset that remained susceptible by conventional MIC criteria. Thus, MIC-defined susceptibility can mask clinically relevant survival heterogeneity that may influence subsequent resistance acquisition and evolution.

This clinical tolerance phenotype differed from the extended-lag or slow-growth tolerance commonly described in laboratory-evolved models. Prolonged lag time can protect bacterial populations during intermittent antibiotic exposure and accelerate resistance evolution^17,19^, yet most tolerant isolates in our collection did not show an extended lag phase or reduced growth rate. These observations indicate that clinical ertapenem tolerance in *K. pneumoniae* is not fully explained by canonical lag-time or slow-growth mechanisms. The identification of *uhpABC* as a candidate tolerance-associated locus, together with the increased ertapenem survival conferred by coordinated expression of the complete operon, provides a functional entry point into the regulatory basis of this clinical survival phenotype.

The temporal and antibiotic-specific patterns further suggest that tolerance was not merely a direct response to ertapenem exposure. Tolerant isolates were already present before ertapenem was introduced locally, and survival under ertapenem correlated with survival under ceftazidime–avibactam, suggesting a β-lactam-associated survival phenotype rather than an ertapenem-specific response. Given the extensive clinical use of β-lactam antibiotics over this period^35,36^, broader β-lactam selective pressures may therefore have contributed to the emergence or persistence of this phenotype. By contrast, cross-tolerance under non-β-lactam antibiotics was restricted to a limited subset of isolates, arguing against a generalized multidrug tolerance.

The enrichment of tolerance among resistant isolates raised an important mechanistic question: whether resistance plasmids directly enhance survival, or whether tolerant recipient backgrounds are more permissive for plasmid acquisition. Our data support the latter interpretation. Plasmid curing reversed high-level ertapenem resistance but did not eliminate ceftazidime–avibactam survival, indicating that resistance plasmids were major contributors to MIC elevation but did not directly confer the tolerant phenotype. Moreover, distinct *bla*_KPC-2_-bearing plasmids among closely related R/T isolates further support repeated plasmid acquisition within tolerant backgrounds rather than expansion after a single plasmid-acquisition event. Together, these findings separate tolerance from resistance plasmid carriage and position tolerance as a recipient-background phenotype that can shape plasmid-mediated resistance evolution.

Conjugation and laboratory evolution experiments linked plasmid-independent tolerance to plasmid-mediated resistance evolution. Tolerant recipient backgrounds showed increased resistance-plasmid acquisition even without antibiotic selection and maintained a larger viable recipient pool during antibiotic exposure, thereby increasing opportunities for plasmid acquisition and persistence under selection. This effect extended beyond initial plasmid acquisition: among carbapenem-resistant isolates that were initially susceptible to ceftazidime– avibactam but differed in pre-existing tolerance to this drug, tolerant backgrounds evolved elevated MICs more rapidly under repeated exposure. Together, these findings suggest that tolerance can shape resistance evolution across successive stages, from plasmid-mediated carbapenem resistance acquisition to later adaptation under ceftazidime–avibactam selection.

Several limitations should be considered. Although the longitudinal collection allowed us to examine tolerance and resistance over more than two decades, the study remains observational at the population scale and cannot prove that each resistant isolate evolved directly from a tolerant ancestor. In addition, tolerance in resistant isolates was assessed using ceftazidime–avibactam survival because high ertapenem MICs made standardized ertapenem killing assays impractical in resistant backgrounds, meaning that susceptible and resistant groups were assayed using different antibiotic readouts. Finally, the mechanisms linking tolerant backgrounds to plasmid acquisition remain incompletely defined. Coordinated *uhpABC* expression increased ertapenem survival, but its necessity across diverse clinical backgrounds and the pathways connecting tolerance to conjugation permissiveness, recipient survival and plasmid establishment require further study.

Together, our study identifies clinical antibiotic tolerance as a pre-existing, MIC-hidden survival phenotype in *K. pneumoniae* that is largely not explained by canonical lag-time or slow-growth tolerance. By linking this phenotype to resistance-plasmid acquisition, recipient survival under antibiotic pressure and subsequent resistance adaptation, our findings broaden current models of the tolerance–resistance relationship from mutation-centered pathways to plasmid-mediated evolution. Integrating survival-based phenotyping with genomic surveillance may help identify tolerant *K. pneumoniae* populations with increased potential for high-level carbapenem resistance emergence.

## Materials and Methods

### Clinical isolate collection

A total of 853 clinical *Klebsiella pneumoniae* isolates collected between 1997 and 2020 were recovered from normally sterile clinical specimens, including blood, cerebrospinal fluid and pleural fluid, at Peking Union Medical College Hospital. In addition, an independent collection of 51 NDM-producing carbapenem-resistant *K. pneumoniae* (NDM-CRKP) isolates was obtained from the Second Affiliated Hospital, Zhejiang University School of Medicine. NDM-CRKP status was defined by carbapenem resistance and the presence of *bla*_NDM_, as determined by genome-based resistance gene screening described below. Unless otherwise stated, longitudinal analyses were performed using the Peking Union Medical College Hospital cohort, whereas the Zhejiang University collection was analyzed as an additional NDM-CRKP cohort.

The use of clinical isolates from Peking Union Medical College Hospital was reviewed and approved by the Ethics Committee of Peking Union Medical College Hospital (approval no. I-22PJ396). The use of NDM-CRKP isolates from the Second Affiliated Hospital, Zhejiang University School of Medicine, was reviewed and approved by the ethics committee of the Second Affiliated Hospital, Zhejiang University School of Medicine (approval no. 2026-0307). All isolates were analyzed retrospectively after routine clinical microbiological testing, and associated clinical information was de-identified before analysis.

Species identification was performed using matrix-assisted laser desorption/ionization time-of-flight mass spectrometry (MALDI–TOF MS). A score of ≥ 2.0 was considered species-level identification according to the recommended interpretation criteria for the Bruker MALDI Biotyper system^37^. All isolates were stored at −80 °C until subsequent phenotypic characterization.

### Minimum inhibitory concentration assays

MICs were measured using a modified broth microdilution assay in LBL medium, adapted from standard broth microdilution approaches for MIC determination^38,39^. Briefly, bacteria recovered from frozen stocks were grown in LBL medium to exponential phase and diluted in fresh LBL medium to a final inoculum of approximately 5 × 10^5^ CFU ml^-1^. Bacterial suspensions were dispensed into 96-well microtiter plates containing two-fold serial dilutions of the indicated antibiotics. Plates were incubated at 37 °C for 24 h, and MICs were recorded as the lowest antibiotic concentration that prevented visible bacterial growth. For β-lactam/β-lactamase inhibitor combinations, avibactam was maintained at a fixed concentration of 4 μg ml^-1^, whereas the partner β-lactam was serially diluted^40^.

### Antibiotic killing assays

Antibiotic killing assays were performed using a modified time-kill assay^11,17^. Isolates were revived from frozen stocks and cultured in LBL medium for 24 h at 37 °C with shaking to obtain stationary-phase cultures. Approximately 10^7^ CFU from each culture were transferred into fresh LBL medium containing the indicated antibiotic at 30× the isolate-specific MIC, unless otherwise stated. For the primary ertapenem tolerance screen, cultures were exposed to ertapenem for 6 h. Killing assays with ceftazidime–avibactam, aztreonam–avibactam, kanamycin and polymyxin E were performed using the antibiotic-specific exposure conditions listed in Supplementary Table 2.

Aliquots were collected before and after antibiotic exposure, serially diluted in sterile phosphate-buffered saline and plated on antibiotic-free LBL agar. Colonies were counted after incubation at 37 °C, and survival fractions were calculated as CFU after treatment divided by CFU before treatment. For the primary ertapenem screen, isolates with survival fractions at least tenfold higher than that of *K. pneumoniae* ATCC 43816 under the same assay conditions were classified as tolerant. All assays were performed in biological triplicate unless otherwise stated.

### ScanLag analysis

Colony appearance-time distributions were quantified using ScanLag, as described previously with minor modifications^41,42^. Isolates were revived from frozen stocks, cultured in LBL medium for 24 h at 37 °C with shaking, serially diluted in sterile phosphate-buffered saline and plated on antibiotic-free LBL agar to obtain well-separated colonies. Plates were incubated at 37 °C and imaged automatically at 20-min intervals for 48 h. Colony appearance time was defined as the time at which an individual colony first exceeded the detection threshold during time-lapse imaging. Appearance-time distributions were extracted using the ScanLag analysis pipeline and compared with those of *K. pneumoniae* ATCC 43816 analyzed under the same conditions. Isolates with an appearance-time delay of ≥600 min relative to ATCC 43816 were classified as having an extended colony-appearance-time phenotype.

### Growth curve

Growth curves were measured in LBL medium using a microplate-reader-based assay^43^. Isolates were revived from frozen stocks, cultured in LBL medium for 24 h at 37 °C with shaking and diluted in fresh LBL medium to approximately 10^5^ CFU per well. Bacterial suspensions were dispensed into 96-well microtiter plates, with medium-only wells included for background correction. Plates were incubated at 37 °C with shaking in a microplate reader, and absorbance at 600 nm was recorded every 10 min for 24 h. Growth curves were generated from background-corrected absorbance values. Growth rates were calculated as the slope of log_10_-transformed absorbance values during the exponential-growth phase by linear regression. The same exponential-phase selection criteria were applied across isolates. All assays were performed in biological triplicate unless otherwise stated.

### Whole-genome sequencing, assembly and annotation

Of the 853 clinical *Klebsiella pneumoniae* isolates, 252 were selected for whole-genome sequencing based on representative phenotypes. Genomic DNA was extracted using the Bacterial Genome DNA Kit (DP302, Tiangen Biotech) according to the manufacturer’s instructions. Sequencing libraries were generated and sequenced on an Illumina NovaSeq 6000 platform to produce 150-bp paired-end reads with a minimum raw-read depth of 100×. Raw reads were quality filtered using Trimmomatic to remove adapter sequences and low-quality bases before assembly. Short reads were assembled de novo using SPAdes v4.1.0^44^. Genome annotation was performed using Prokka v1.14.5^45^. Sequence types, capsular and O-antigen locus types, and carbapenemase genes were assigned using Kleborate v3.2.4^46^. Carbapenemase gene calls were confirmed by BLASTn against curated reference sequences from the NCBI nucleotide database. Plasmid-associated contigs, replicon types, mobility predictions and plasmid cluster assignments were determined using MOB-suite v3.1.9^47^. Plasmid clusters were interpreted as MOB-suite-defined whole-sequence-based plasmid groups. Carbapenemase genes were considered plasmid-associated when the corresponding gene-containing contig was assigned to a MOB-suite plasmid reconstruction or carried plasmid-associated features.

### Core-genome cluster network and reference-based phylogenetic analysis

A core-genome-based isolate-relatedness network was generated to visualize genetically related clinical *K. pneumoniae* isolates. Only connected isolates were retained for network visualization. Pairwise core-genome distances were calculated from the core-genome alignment and used to construct an undirected network connecting genetically related isolates. The network was visualized in Gephi v0.10, with node metadata annotated by sequence type, resistance phenotype and tolerance phenotype^48^.

Reference-based phylogenetic reconstruction was performed for the 252 sequenced *K. pneumoniae* isolates. Short reads were mapped to the complete chromosome assembly of the clinical isolate K13, generated by long-read sequencing in this study, using Snippy v4.6.0 to generate a core-genome SNP alignment. Recombinant regions were identified and masked using Gubbins v2.4.1^49^, and a maximum-likelihood tree was inferred from the recombination-filtered SNP alignment using IQ-TREE v3.0.1^50^. The tree was visualized using iTOL^51^.

### Plasmid curing

Plasmid curing was performed using plumbagin, adapted from established plasmid-curing approaches^33^. Briefly, approximately 10⁷ CFU of each selected *bla*_KPC-2_-positive *K. pneumoniae* isolate were inoculated into LBL medium supplemented with 50 μg ml^-1^ plumbagin and incubated overnight at 37 °C with shaking at 220 r.p.m. Cultures were serially diluted and plated on antibiotic-free LB agar to obtain single colonies. Candidate cured derivatives were screened by PCR for loss of *bla*_KPC-2_. PCR-negative colonies were further tested for ertapenem MICs using the modified broth microdilution assay described above. Colonies showing loss of *bla*_KPC-2_ together with a marked reduction in ertapenem MIC were considered derivatives lacking the carbapenemase-encoding plasmid and were used for phenotypic assays as indicated.

### In vitro conjugation experiment

In vitro conjugation assays were performed using a filter-mating approach^52,53^. *Escherichia coli* BW25113 carrying the *bla*_NDM_-positive plasmid p3R-4 was used as the donor, and clinical *Klebsiella pneumoniae* isolates were used as recipients. Recipient isolates were pre-screened for growth on LB agar containing kanamycin (50 μg ml^-1^). Donor cultures were grown in selective LB broth, diluted 1:100 into fresh selective medium and cultured to logarithmic phase. Recipients were inoculated from single colonies and cultured in LB broth for 24 h at 37 °C with shaking.

Donor and recipient cells were washed twice with sterile phosphate-buffered saline, mixed at a donor: recipient ratio of 1:3 and spotted onto sterile 0.22-μm hydrophilic membrane filters placed on antibiotic-free LB agar. After incubation at 37 °C for 3 h, bacteria were recovered from the filters, serially diluted and plated on LB agar containing ceftazidime–avibactam (ceftazidime, 20 μg ml^-1^; avibactam, 4 μg ml^-1^) and kanamycin (50 μg ml^-1^) to select transconjugants. Selection specificity was verified using donor-only and recipient-only controls. Conjugation frequency was calculated as transconjugant CFU divided by recipient CFU. Experiments were performed in biological triplicate.

### Cyclic evolution protocol

Cyclic evolution experiments were performed using a repeated ceftazidime–avibactam exposure, wash and regrowth protocol^17^. Bacterial populations recovered from frozen stocks were cultured in LBL medium for 24 h at 37 °C with shaking at 220 r.p.m. Cultures were then diluted 1:100 into 50 ml fresh LBL medium containing ceftazidime–avibactam at 10× the baseline ceftazidime–avibactam MIC of each starting isolate, with avibactam maintained at 4 μg ml^-^¹, and incubated for 6 h at 37 °C with shaking at 220 r.p.m.

After antibiotic exposure, cells were collected by centrifugation at 6,000g for 15 min and washed three times with 1 ml sterile phosphate-buffered saline to remove residual antibiotic. The final cell pellet was resuspended in 1 ml fresh LBL medium and split into two aliquots. One aliquot was cultured for 24 h at 37 °C with shaking at 220 r.p.m. to initiate the next cycle, whereas the other aliquot was stored as a frozen stock and used for phenotypic analysis of the corresponding cycle. This antibiotic-exposure–wash–regrowth procedure was repeated for 5 cycles. MICs were determined at the indicated cycles using the modified broth microdilution assay described above.

### Phylogeny-guided identification, variant enrichment and functional validation of the *uhpABC* locus

To identify candidate genetic determinants associated with ertapenem tolerance while reducing confounding by broad population structure, we first selected a closely related lineage from the core-genome relatedness analysis that contained both ertapenem-susceptible tolerant and ertapenem-susceptible non-tolerant isolates. This analysis nominated the *uhpABC* locus as a candidate tolerance-associated region. Targeted amino-acid variant enrichment across the *uhpABC* locus was performed using predicted UhpC, UhpB and UhpA protein sequences from sequenced clinical isolates. Reference sequences were excluded from enrichment testing. Because UhpB contained a 2-aa insertion/deletion polymorphism, UhpB sequences were globally aligned and converted to a unified 494-aa coordinate system. UhpC, UhpB and UhpA were concatenated by isolate to generate a 1,135-aa operon-level alignment. Variants relative to the reference sequence were tested for phenotype enrichment using Fisher’s exact test, with Benjamini–Hochberg false-discovery-rate correction. A separate tolerant-versus-susceptible comparison was performed after excluding resistant isolates. Enriched variants were mapped onto the annotated *uhpABC* locus, and UhpB domain boundaries were assigned using EBI InterPro annotation^54^. The *uhpABC* locus was annotated based on homology to the canonical UhpABC sugar-phosphate regulatory system^34^.

For functional validation, *uhpA*, *uhpB*, *uhpC* and the complete *uhpABC* operon were amplified from K997 and cloned into the IPTG-inducible pET-ECFP vector by replacing the ECFP coding sequence. The original ECFP-expressing vector was used as the control. Sequence-confirmed constructs were introduced into *K. pneumoniae* ATCC 43816, induced with 1 mM IPTG, and tested for ertapenem survival using the killing assay described above. Survival fractions were calculated as CFU after ertapenem exposure divided by CFU before exposure. All assays were performed in biological triplicate unless otherwise stated.

### Statistical analyses and data visualization

Statistical analyses and data visualization were performed using GraphPad Prism version 10.1.1 and R version 4.4.2. Survival fractions were calculated as viable counts after antibiotic exposure divided by viable counts before treatment. Survival rates, MIC values and conjugation efficiencies were analyzed on a logarithmic scale where appropriate. Statistical tests, definitions of n, center values, error bars and exact P values are provided in the corresponding figure legends. Unless otherwise stated, statistical tests were two-sided, *P* values <0.05 were considered statistically significant, and multiple testing was controlled using the Benjamini–Hochberg false-discovery rate procedure where applicable. Final figures were assembled in Adobe Illustrator.

### Supplementary Materials

The PDF file includes:

Extended Data Fig. 1 to 6

Extended Data Table 1

Other Supplementary Material for this manuscript includes the following:

Supplemental Tables 1-2

## Acknowledgement

We thank Jianzhong Shen (College of Veterinary Medicine, China Agricultural University) for providing *Escherichia coli* BW25113/p3R-4 and Jingren Zhang (School of Basic Medical Sciences, Tsinghua University) for providing *Klebsiella pneumoniae* ATCC 43816. We are grateful to Wei Kang and Yanbing Li (Department of Clinical Laboratory, Peking Union Medical College Hospital) for assistance with clinical isolate collection, and to Junhao Zhu (Institute of Microbiology, Chinese Academy of Sciences), Jintao Liu and Guanxiang Liang (both School of Basic Medical Sciences, Tsinghua University) for helpful discussions and constructive suggestions during this study. This study was funded by the National Key Research and Development Program of China (2022YFC2303202), the Tsinghua-Peking Joint Center for Life Sciences (20111770319), and the Tsinghua University Dushi Program (20261080009) to J. L.

## Author Contributions

W.Z. designed, performed experiments, analyzed the data and drafted the manuscript. B.Z. designed, performed experiments, conducted the bioinformatic analysis and edited the manuscript. M.Z., Y.X., and R.Z. participated in study design, strain collection and edited the manuscript. J.L. supervised the whole project and edited the manuscript.

## Competing Interests

The authors declare no competing interests.

## Data and Materials Availability

All data needed to evaluate the conclusions in the paper are present in the paper and/or the Supplementary Materials

## Code availability

No custom software or algorithms were developed for this study. Statistical analyses and figure generation were performed in R using standard packages, as described in the Methods. The R scripts used to generate the figures are available from the corresponding author upon reasonable request.

## Notes

### Competing Interest Statement

The authors have declared no competing interest.

