## Supplementary Information for "Pre-existing antibiotic tolerance facilitates plasmid-mediated carbapenem resistance evolution in clinical *Klebsiella pneumoniae*"

**The PDF file includes:**

Extended Data Fig. 1 to 6

Extended Data Table 1

**Other Supplementary Material for this manuscript includes the following:**

Supplemental Tables 1-2

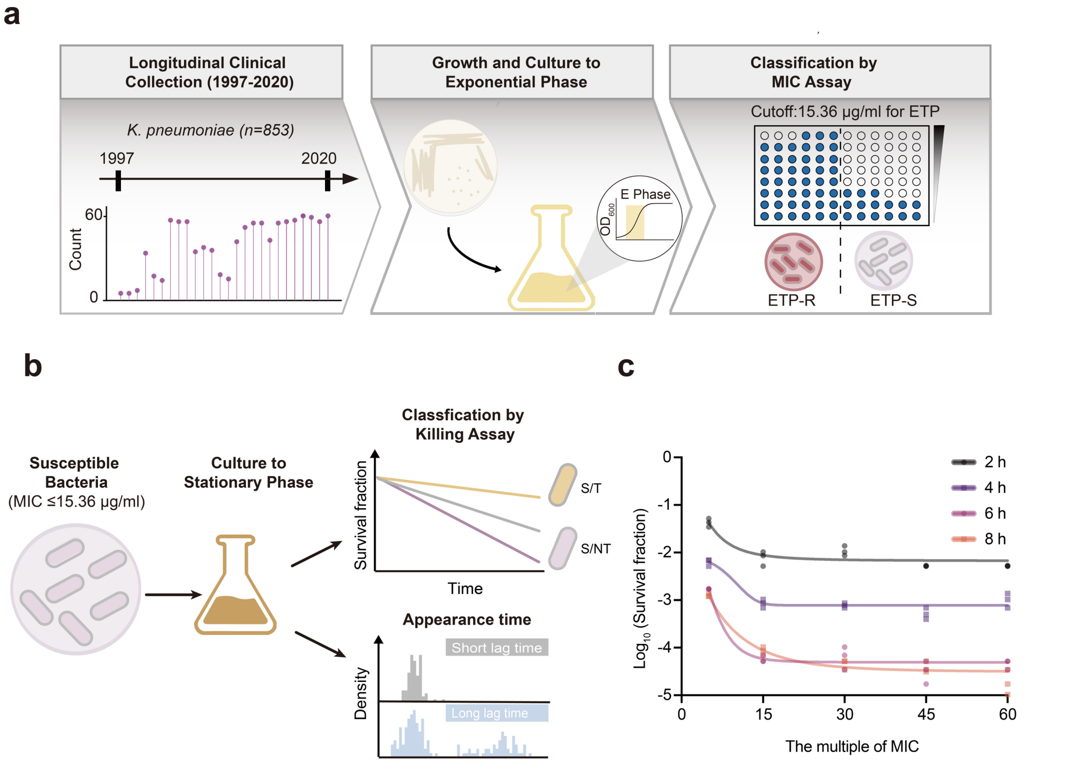


**Extended Data Fig. 1 | Workflow for antibiotic tolerance screening in clinical *Klebsiella pneumoniae* isolates.** **a**, Retrospective assembly and MIC-based stratification of the longitudinal clinical *K. pneumoniae* cohort. A total of 853 clinical *K. pneumoniae* isolates collected between 1997 and 2020 were analyzed. Individual isolates were revived and cultured to exponential phase before antimicrobial susceptibility testing. Using a study-defined cutoff of 15.36 μg ml$^{-1}$, isolates were classified as ertapenem-resistant (ETP-R) or ertapenem-susceptible (ETP-S). **b**, Schematic of the workflow used to quantify antibiotic tolerance among ertapenem-susceptible isolates. Isolates were cultured to stationary phase and subjected to ertapenem killing assays to classify MIC-defined susceptible isolates into tolerant (S/T) and non-tolerant (S/NT) populations. Colony appearance times were measured using ScanLag to assess lag-phase heterogeneity. **c**, killing assay optimization using the reference strain *K. pneumoniae* ATCC 43816. Survival was measured after exposure to ertapenem at different multiples of the isolate-specific MIC for 2, 4, 6 or 8 h. Data are presented as mean ± s.d. from at least three biological replicates.

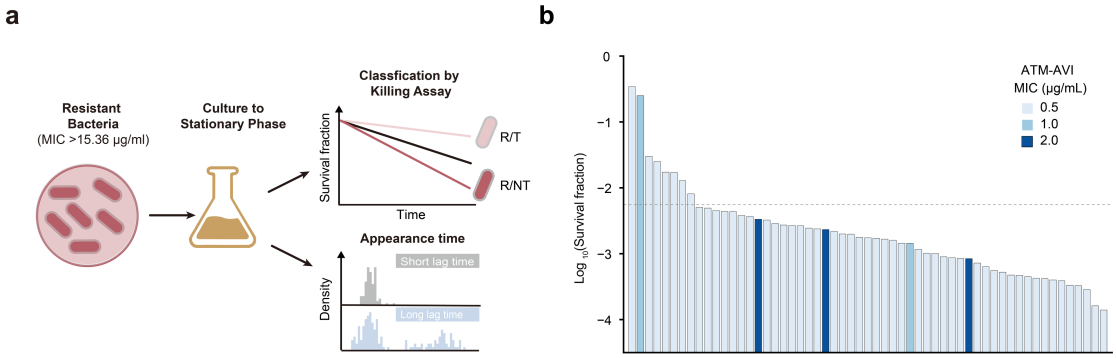


**Extended Data Fig. 2 | Survival heterogeneity under aztreonam–avibactam exposure among NDM-producing CRKP isolates**. **a**, Schematic of the workflow used to quantify antibiotic tolerance among ertapenem-resistant isolates. Isolates with ertapenem MICs >15.36 μg ml-1 were cultured to stationary phase and subjected to killing assays to classify resistant isolates into tolerant (R/T) and non-tolerant (R/NT) populations. Colony appearance times were measured using ScanLag to assess lag-phase heterogeneity. **b**, Survival fractions of NDM-producing CRKP isolates (n = 51) under aztreonam–avibactam (ATM–AVI) exposure. Each bar represents one isolate, ordered by log10-transformed survival fraction. Bar colors indicate the ATM–AVI MIC of each isolate. The dashed line indicates the predefined survival threshold used to classify tolerant isolates. Tolerant isolates accounted for 15.7% of this group (8/51). Most isolates had low ATM–AVI MICs, whereas survival varied substantially across isolates, revealing survival heterogeneity that was not captured by MIC alone.

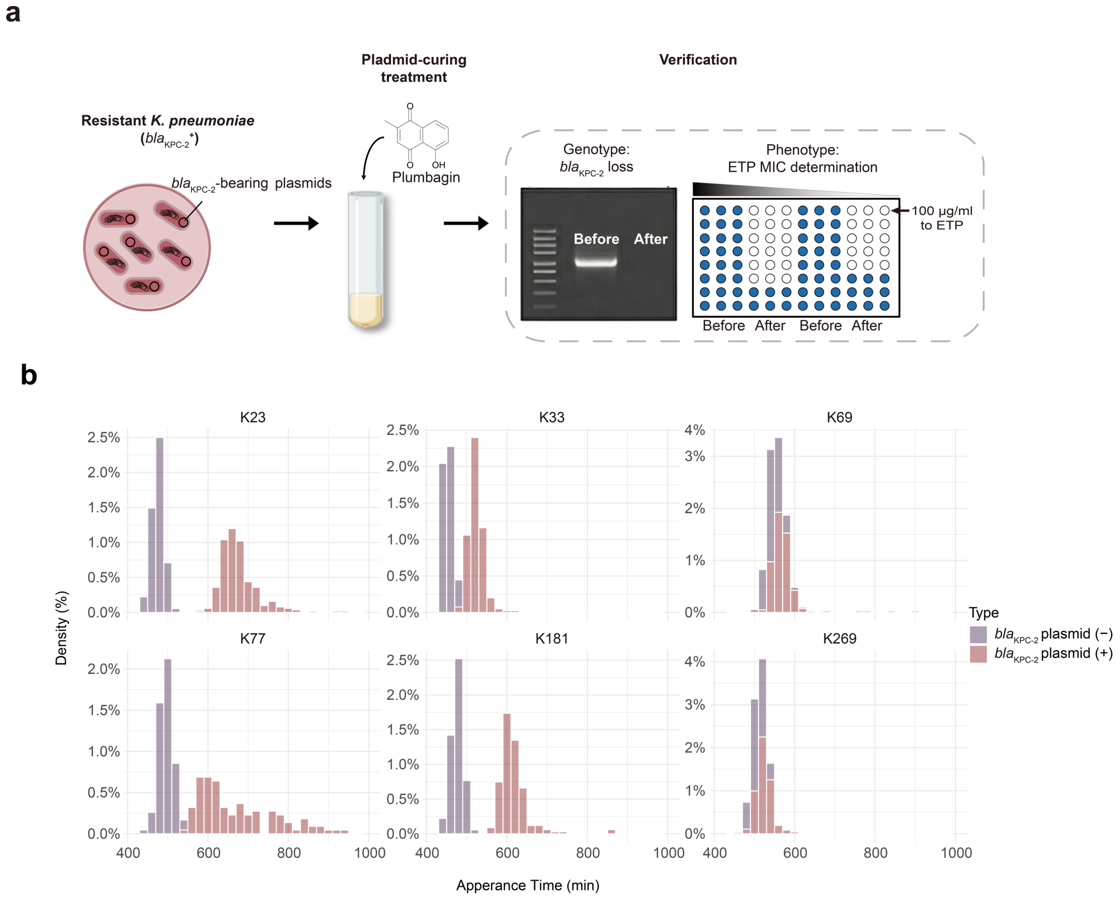


**Extended Data Fig. 3 | Plasmid curing and appearance-time profiling of resistant *K. pneumoniae* isolates**. **a**, Schematic of plasmid curing in high-level ertapenem-resistant *K. pneumoniae* isolates carrying *bla*$_{KPC-2}$-bearing plasmids. Plumbagin-treated isolates were verified by *bla*$_{KPC-2}$ PCR and ertapenem MIC determination. **b**, ScanLag-derived colony appearance-time distributions of six clinical isolates before and after plasmid curing. Plasmid (+) denotes parental plasmid-containing isolates, and plasmid (−) denotes the corresponding plasmid-cured derivatives. The x axis indicates colony appearance time in minutes, and the y axis indicates density.

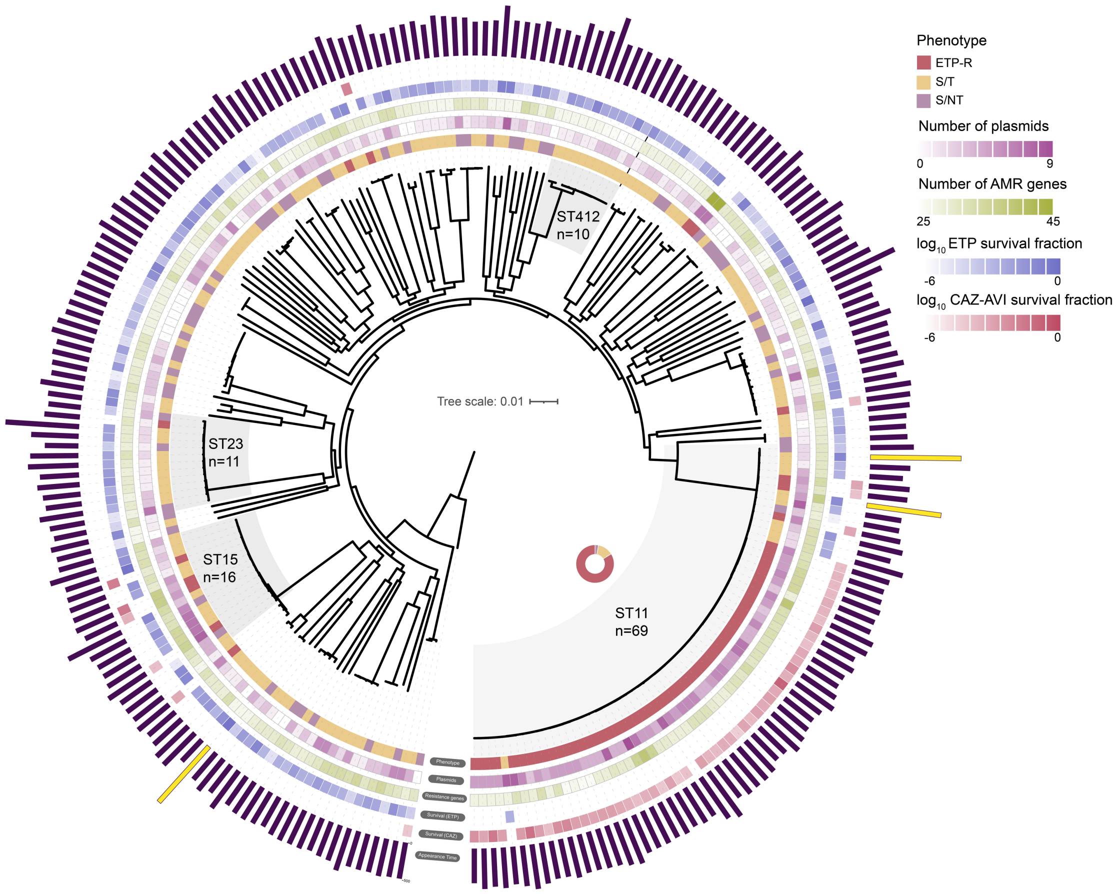


**Extended Data Fig. 4 | Phylogenomic landscape and phenotypic profiling of clinical Klebsiella pneumoniae isolates**. Maximum-likelihood phylogeny of clinical *K. pneumoniae* isolates based on core-genome single-nucleotide polymorphisms (n = 252). Major sequence types are highlighted by shaded sectors, including ST11 (n = 69), ST15 (n = 16), ST23 (n = 11) and ST412 (n = 10). The scale bar indicates substitutions per site. Concentric tracks show, from inner to outer: phenotypic classification, number of plasmids, number of antimicrobial-resistance genes, $\log_{10}$ ertapenem survival fraction and $\log_{10}$ ceftazidime–avibactam survival fraction. Heat maps indicate the values shown in the legend. The outer bar plot shows colony appearance time for each isolate.

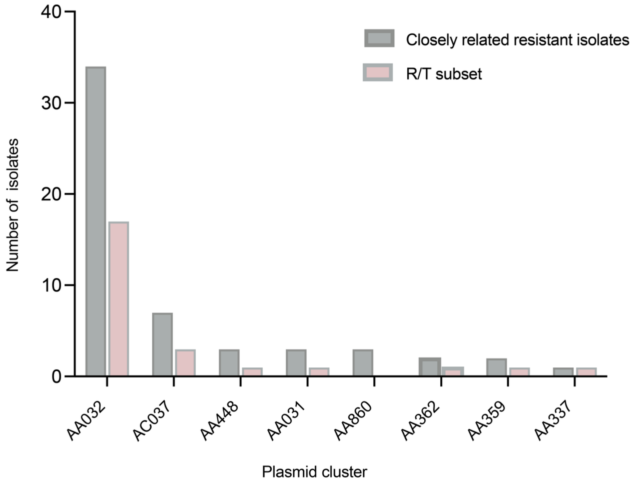


**Extended Data Fig. 5 | Distribution of *bla*$_{KPC-2}$-bearing plasmid clusters among resistant isolates from a closely related lineage.** Bar plot showing the distribution of MOB-suite-assigned *bla*$_{KPC-2}$-bearing plasmid clusters among ertapenem-resistant *K. pneumoniae* isolates within the focal closely related lineage. Grey bars indicate all resistant isolates in this lineage, and pink bars indicate the ertapenem-resistant tolerant subset (R/T). Each bar represents the number of isolates carrying a *bla*$_{KPC-2}$-bearing plasmid assigned to the indicated plasmid cluster. Although AA032 was the dominant plasmid cluster, R/T isolates were distributed across multiple plasmid clusters, consistent with repeated acquisition of resistance plasmids within tolerant backgrounds rather than expansion solely after a single plasmid-acquisition event.

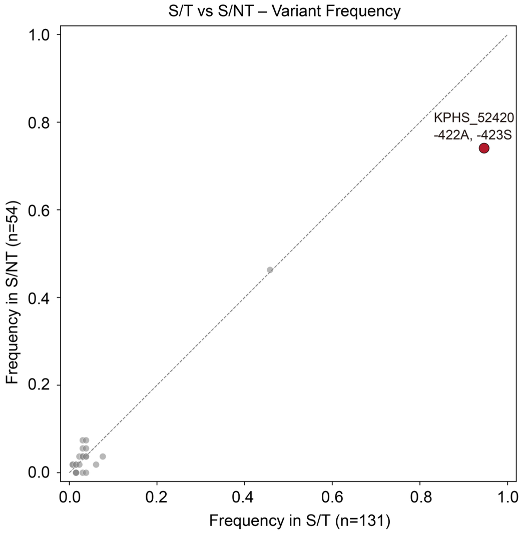


**Extended Data Fig. 6 | Variant-frequency comparison identifies a tolerance-enriched *uhpB* indel within the *uhpABC* locus**. Scatter plot showing the frequency of amino-acid variants within the *uhpABC* locus among ertapenem-susceptible tolerant (S/T; n = 131) and ertapenem-susceptible non-tolerant (S/NT; n = 54) *K. pneumoniae* isolates. Each point represents one variant. The x axis indicates variant frequency in S/T isolates, and the y axis indicates variant frequency in S/NT isolates. The dashed diagonal line indicates equal frequency between the two groups; variants below the diagonal are more frequent in S/T isolates, whereas variants above the diagonal are more frequent in S/NT isolates. The highlighted red point indicates the *uhpB* 422–423 insertion, which was more frequent in S/T isolates but was not exclusive to this group. This cohort-level frequency comparison supports the enrichment signal identified in the within-lineage variant analysis.

**Extended Data Table 1 | Carbapenemase gene composition and plasmid context among sequenced high-level ertapenem-resistant isolates**

| Carbapenemase category | No. of isolates | Percentage | Major ST | Predicted genomic context | Predominant plasmid replicon(s) |
|---|---|---|---|---|---|
| $\mathit{bla}_{\text{KPC-2}}$ | 64/68 | 94.1% | ST11 | Plasmid-associated contigs | IncFIA |
| $\mathit{bla}_{\text{NDM}}$ | 4/68 | 5.9% | ST199 | Plasmid-associated contigs | IncFIB |
| Total | 68/68 | 1 | — | Predominantly plasmid-associated | — |

High-level ertapenem resistance was defined as an ertapenem MIC >100 μg $ml^{-1}$. Data are shown for 68 sequenced high-level ertapenem-resistant isolates. Four ertapenem-resistant isolates had MICs >15.36 and <100 μg $ml^{-1}$; these isolates were retained in analyses of the broader ertapenem-resistant cohort but excluded from analyses specifically referring to high-level ertapenem resistance. Two additional high-level ertapenem-resistant isolates were not sequenced and were therefore excluded from genome-based analysis.
